# Plant-on-a-chip: continuous, soilless electrochemical monitoring of salt uptake and tolerance among different genotypes of tomato

**DOI:** 10.1101/2024.02.03.578647

**Authors:** Philip Coatsworth, Yasin Cotur, Tarek Asfour, Zihao Zhou, José M. R. Flauzino, Tolga Bozkurt, Firat Güder

## Abstract

Tomatoes (*Solanum lycopersicum*), a high-value crop, exhibit a unique relationship with salt, where increased levels of NaCl can enhance flavor, aroma and nutritional quality but can cause oxidative damage and reduce yields. A drive for larger, better-looking tomatoes has reduced the importance of salt sensitivity, a concern considering that the sodium content of agricultural land is increasing over time. Currently, there are no simple ways of comparing salt tolerance between plants, where a holistic approach looking at [Na^+^] throughout the plant typically involves destructive, single time point measurements or expensive imaging techniques. Finding methods that collect rapid information in real time could improve the understanding of salt resistance in the field. Here we investigate the uptake of NaCl by tomatoes using TETRIS (**T**ime-resolved **E**lectrochemical **T**echnology for plant **R**oot environment ***I****n-situ* chemical **S**ensing), a platform used to measure chemical signals in the root area of living plants. Low-cost, screen-printed electrochemical sensors were used to measure changes in salt concentration *via* electrical impedance measurements, facilitating the monitoring of the uptake of ions by roots. We not only demonstrated differences in the rate of uptake of NaCl between tomato seedlings under different growth conditions, but also showed differences in uptake between varieties of tomato with different NaCl sensitivities and the relatively salt-resistant “wild tomato” (*Solanum pimpinellifolium*) sister species. Our results suggest that TETRIS could be used to ascertain physiological traits of salt resistance found in adult plants but at the seedling stage of growth. This extrapolation, and the possibility to multiplex and change sensor configuration, could enable high-throughput screening of many hundreds or thousands of mutants or varieties.

## Introduction

NaCl, or simply “salt”, is toxic for the majority of plants, leading to oxidative damage and reduced crop yields.^1, 2^ For tomatoes, however, there is a push and pull with salt: although more salt leads to lower yields and smaller fruits, those fruits will often have better flavor, aroma and nutritional properties, where increasing salinity increases concentrations of fructose, glucose, minerals, carotene and vitamin C.^3, 4^ Tomatoes (*Solanum lycopersicum*) are a valuable crop plant, with US production having a value of around US $1.8 billion in 2022 – “heirloom” varieties are sold at a higher premium due to their superior taste and quality.^5, 6^ Unfortunately, a drive for larger, more attractive fruits has reduced the priority of organoleptic properties, nutrition and salt tolerance during tomato production, a concern considering that the sodium content of agricultural land is increasing over time.^7^ Breeding with other species closely related to tomato (such as the “wild tomato” species *Solanum pimpinellifolium*) could increase salt and disease resistance in cultivated tomatoes, while improving their nutritional quality.^8^

In agriculture, electrical conductivity (EC) measurements are used to ascertain the overall ionic activity of a nutrient solution or growth medium, which can give an indication into how the crop will develop.^3^ Generally, high EC will produce tomato fruits that are highly flavored, but with reduced fruit sizes and yield. Excess salinity can decrease water availability to the plant due to reduced osmotic pressure across the roots, which in turn reduces xylem transport of water and solutes from the roots to the fruits. The specific ionic content of the medium is also important to consider, where high EC water that contains an excess of Na^+^ ions may prevent uptake of macronutrients, such as K^+^, due to competing transport.^9^

In some countries, the volume and quality of water used in irrigation may be reduced, leading to higher EC values and therefore lower yield.^10^ In other areas, tomatoes are often grown covered in greenhouses or under plastic-covered tunnels to prevent frost damage, and where soilless systems are used, the reuse of irrigation water may reduce costs but can lead to increasing EC of the water over time.^3, 5^ Across the world, arable land is also vastly outsized by marginal land (land that is unsuitable for farming), where salt-affected land forms a large proportion of marginal land.^11^ In these high-salinity conditions, the use of cultivars with increased salt resistance would be beneficial and could potentially recover these marginal lands for agricultural use. While varieties exist with a range of sensitivities to salt, *S. lycopersicum* is generally not considered resistant to NaCl. It has been proposed that wild tomato genotypes, such as *S. pimpinellifolium*, may be good candidates for gene donation to improve salt tolerance in commercial cultivars.^8^

Beyond its purely agricultural value, *S. lycopersicum* is also of great importance to the biological field as a model plant. The tomato has features (including flesh fruits and compounds leaves) not present in other model plants, such as O*ryza sativa* (rice) and *Arabidopsis thaliana*. *S. lycopersicum* is also closely related to other members of the *Solanaceae* family, which includes potato, eggplant, tobacco, and is studied for its response to biotic (pathogenic) and abiotic (such as light, salinity and temperature) stresses.^12, 13^ Beyond being an essential part of photosynthesis, light is known to affect many plant processes, including growth, nutrient uptake and salt tolerance.^14, 15^

Currently, there are no simple ways of measuring or comparing salt tolerance or sensitivity between plants, and instead a holistic approach must be taken. When looking at differences in salt tolerance between tomato genotypes, researchers often measure variation in Na^+^ transport from outside the root into the shoot, correlation between concentration of Na^+^ and leaf area, Na^+^ accumulation in old leaves compared to new, and leaf [K^+^]/[Na^+^] ratio.^16, 17^ Typically, these four “physiological characters” may only be established destructively, relying on dried plant mass or by taking physical samples, and single measurement cannot capture complex, time-dependent plant behavior. As these characteristics could be useful for plant breeding programs, finding methods that collect rapid information in real time could improve the understanding of salt resistance in the field. Salt stress can produce visual and physical symptoms that range in severity, such as yellowing, leaf drying and rolling, tip whitening, wilting, premature senescence, cessation of growth and death.^18^ These measurements are often not quantitative, however, and there is a need for quantitative methods and technology that can relieve the current “bottleneck” in phenotyping of different (engineered) varieties.^19^ Imaging methods can provide rich time-dependent information, such as positron emission tomography (PET), which has been used to monitor the transport of ^22^Na^+^ in various plant species.^20, 21^ These methods often require expensive specialized equipment, however. The quality of data can also depend on optimum setup and optical inference.^22^

In this work, we share our research into the uptake of salt by tomato plants, utilizing our whole-plant, soilless monitoring platform TETRIS (**T**ime-resolved **E**lectrochemical **T**echnology for plant **R**oot ***I****n-situ* chemical **S**ensing). TETRIS comprises low-cost, 2D screen-printed electrochemical sensors located underneath the roots of living plants to measure the local chemical environment. Due to the importance of sodium by tomatoes for both the quality of the fruits and for salt sensitivity, we used TETRIS to monitor the uptake of NaCl in *S. lycopersicum via* electrochemical impedance measurements. As light is another key factor in the growth of plants and uptake of ions, we also investigated the uptake of NaCl and KNO_3_ in plants grown under dark conditions compared to a standard light-dark schedule, where plants grown in dark generally showed higher uptake. Finally, as different varieties of tomatoes and related species can have differences in their tolerance to salt, we have utilized TETRIS to elucidate differences in Na^+^ uptake between commercial cultivars of *S. lycopersicum* and compared these to the salt-resistant wild tomato species *S. pimpinellifolium*. By demonstrating higher uptake of NaCl in sensitive plants, we demonstrate TETRIS could be used to show salt resistance of mature plants at the seedling stage of growth.

## Results and discussion

### General experimental setup of TETRIS

TETRIS consisted of a measurement chamber and a disposable sensing module (**Figure 1A**).^23^ The chamber comprised a transparent acrylic lid and silicone base with water reservoir, allowing the plant to photosynthesize whilst maintaining a constant humidity. Screen-printed electrodes on polyester transparency sheet were affixed to a raised platform, placed into the measurement chamber and connected to a potentiostat (PalmSens 4 by PalmSens BV, Netherlands). The use of two eight-channel multiplexers allowed for the simultaneous monitoring of up to 16 sensors. To detect the chemical environment around the roots of living plants, all experiments were performed where plants were grown on chromatography paper and placed onto the sensing module. Firstly, seeds were washed and germinated on wet tissue before transferring onto discs of chromatography paper. The area of growth of the chromatography paper was defined by hydrophobic wax barriers formed by a HPRT MT800 thermal transfer printer. The discs with germinated seeds were then placed in enclosed boxes and supplied with water from a reservoir for a set number of days, before being removed from the box and placed onto the sensing module to perform experiments.

**Figure 1.**
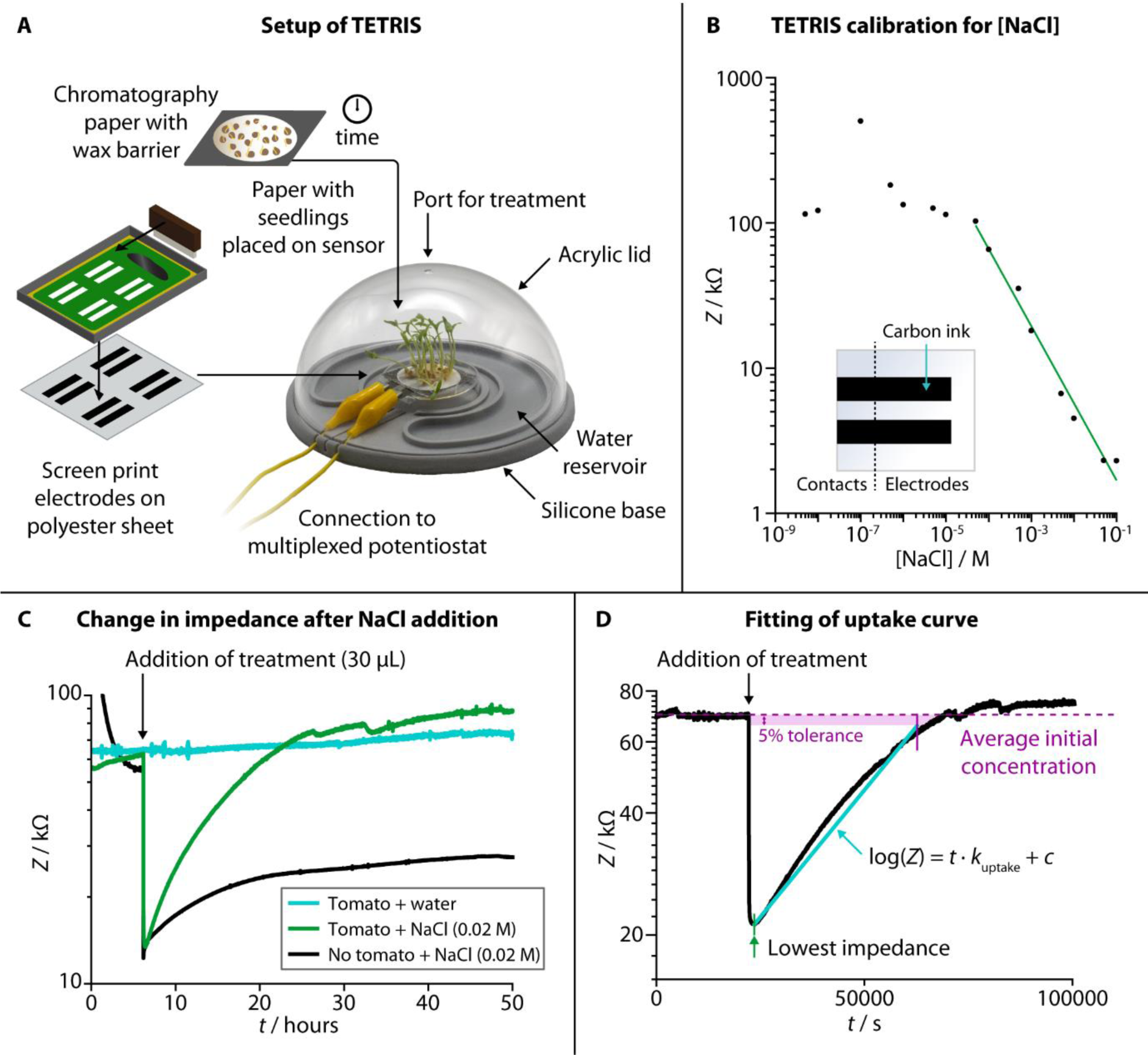
Setup, characterization and experimental use of TETRIS. (**A**) Schematic showing the setup of TETRIS, including production of sensors and preparation of seedlings. (**B**) Calibration of the impedance sensor in TETRIS for [NaCl], with insert showing sensor design. (**C**) By measuring the impedance of a filter paper disc with tomato seedlings (“Heinz 1350”, 20 plants, 7 days growth, green signal), we could monitor the uptake of NaCl upon addition, compared to filter paper disc with no seedlings (black signal) or paper disc with seedlings, but addition of deionized water (cyan signal). (**D**) The rate of uptake of NaCl by tomato seedlings (“Heinz 1350”, 20 plants, 7 days growth) could be found by plotting log(*Z*) against time and fitting a line from the point of lowest impedance after addition of treatment and the point at which the impedance returns to the initial average value (if at all), with 5% tolerance. The rate of uptake, *k*_uptake_, was set as the gradient of the line.

We used two conductive printed carbon electrodes to measure electrical impedance of the solution in the paper disc between the two electrodes. As electrical impedance decreases with increased salt concentration due to higher number of charge carriers (**Figure 1B**, NaCl calibration curve), we have used this sensing setup to monitor the uptake of ions by living plants.

When a disc of chromatography paper wetted with deionized water was placed on the impedance sensor, the initial electrical impedance was relatively high (>50 kΩ) due to a lack of ions. Upon addition of NaCl solution (**Figure 1C**), the impedance dropped (to around 10 – 20 kΩ) due to an increase in the number of charge carries in the paper. An increase in impedance (around 5 – 10 kΩ) occurred over tens of hours due to the added ions diffusing through the solution in the paper, but it remained lower than the initial baseline impedance (before salt was added). When living plants were present on the paper discs, the impedance dropped when salt was added by a similar amount, but, over tens of hours, the impedance steadily rose back up to the baseline level. This effect is attributed to the uptake of ions from the solution in the paper by the seedlings present on top. We have previously observed a similar effect in kale seedlings for multiple salts with a range of ions (including sodium salts, heavy metal salts and nutrients), and TETRIS is not limited to just measuring changes in [NaCl].^23^ To compare this observed uptake between experiments, the rate of uptake, *k*_uptake_, was calculated by taking the gradient of the slope in the log(*Z*) *vs. t* plot (**Figure 1D**).

### Continuous monitoring of uptake of ions in the root environment of tomato seedlings

We used TETRIS to measure the uptake of added NaCl by tomato seedlings with different experimental conditions. Initially, we compared uptake between different numbers of tomato seedlings (“Heinz 1350”) grown in the same way on paper discs, where we found a greater number of plants showed greater rates of uptake of NaCl (30 µL, 0.02 M) from zero to ten plants (**Figure 2A**). This is unsurprising, due to the greater surface area of roots available to take up ions with a greater number of plants. Greater increase was not observed between ten and twenty plants, presumably due to all the added ions being taken up by ten plants, although twenty plants were used subsequently to ensure high levels of uptake. Weighing the mass of these different numbers of plants also showed that greater rate of uptake corresponded to greater mass, for both fresh mass (**Figure S1A**) and dry mass (**Figure 2B**).

**Figure 2.**
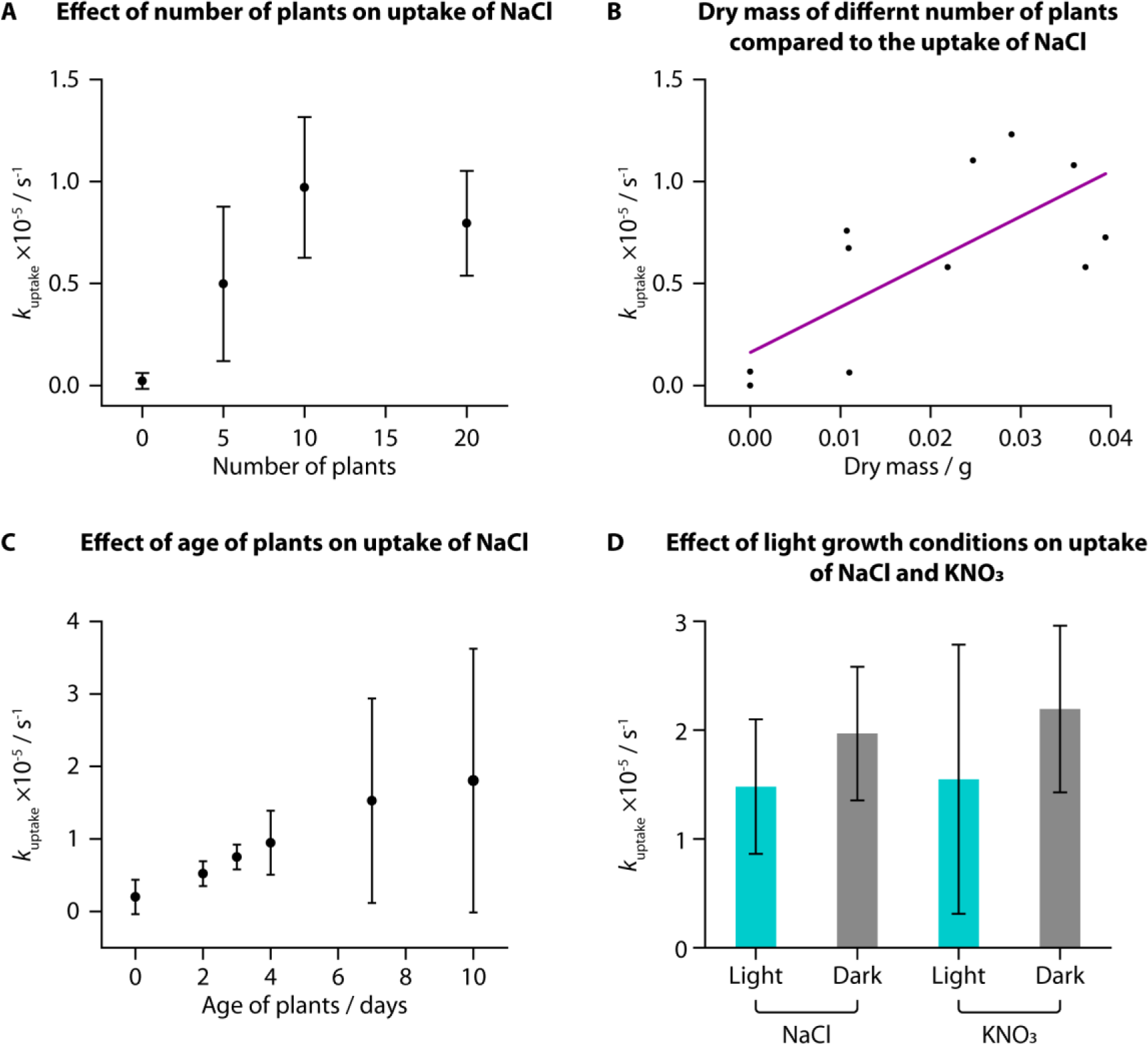
Uptake of ions measured in *S. lycopersicum* “Heinz 1350” under different experimental conditions. (**A**) *k*_uptake_ for NaCl (addition of 30 µL, 0.02 M) generally increased with number of tomato seedlings (7 days post germination: n = 3). This corresponded to dry mass of the seedlings after the experiment (**B**). (**C**) There was an increase in *k*_uptake_ for NaCl (addition of 30 µL, 0.02 M) with age of plants (20 plants), with large deviation in *k*_uptake_ (2 days post germination: n = 6; 3 days: n = 5; all others: n = 3). (**D**) Plants grown in a dark environment, on average, had a larger *k*_uptake_ for both NaCl and KNO_3_ (addition of 30 µL, 0.02 M) than plants grown in a standard light-dark schedule, however this result did not yield statistically significant differences (KNO_3_, dark: n = 4; all others: n = 3). Lighting conditions, therefore, appear not to affect salt or nutrient uptake rates.

When monitoring uptake with seedlings of different ages, we observed an increase in average *k*_uptake_ with increasing age of the plants from 2 days to 10 days post-germination for addition of NaCl (30 µL, 0.02 M). The deviation between samples was large, however, and while the fresh mass of the samples increased with age, the dry mass was not notably different (**Figure S1B-C**). It is likely that the samples did not have large differences in the total root area, despite their difference in ages, resulting in similar uptake amounts.

*S. lycopersicum* has been utilized to study the effects of light stress for both high and low light levels. Low light levels are associated with higher risk of pathogenic infection, and the color and intensity of light can lead to accumulation of specific metabolites.^13^ Steroidal glucoalkaloids, which are toxic to humans, were found to have greater accumulation under red, blue and fluorescent light sources, although the underlying mechanisms are unclear.^24^ High intensity light has been found to induce nonuniform pigmentation patterns of anthocyanins (pigments that can aid with protection from UV and high intensity light), and low intensity light did not activate genes for anthocyanin biosynthesis at all.^25^ Higher light intensities are also linked to higher uptake of nutrients (K and P) in tomato plants and increased biomass.^15^

We grew tomato seedlings as before but placing the growth box in the dark for the entire growth period. We then compared the uptake of NaCl and KNO_3_ (both 30 µL, 0.02 M) to tomatoes grown in the dark compared to those grown in the standard laboratory light-dark schedule (approximately 8h dark and 16h light per day). Interestingly, despite previous research suggesting that greater light leads to higher nutrient uptake, we did not observe a statistically significant difference in uptake of either NaCl or KNO_3_ for plants grown in different lighting conditions, when using a two-sample *T*-test (NaCl: *t*(6) = −1.1209, *p* < 0.05; KNO_3_: *t*(7) = −0.96669, *p* < 0.05). We did, however, find that the plants grown in the dark were notably taller (**Figure S1D**, average height of 5.8 cm) compared to plants grown in standard lighting conditions (average height of 4.5 cm). This difference in height was also found to be statistically significant when using a two-sample *T*-test (*t*(8) = −4.5018, *p* < 0.05). Conversely, however, these dark-grown plants had a lower average fresh and dry mass than those grown in standard lighting conditions and visibly were more yellow and unhealthy (**Figure S1E-G**).

This phenomenon of plants grown from seed in full darkness being taller and more yellow is well documented.^26, 27^ Under a standard light-dark schedule, once germinated seedlings have reached the surface of the soil and are exposed to light, they will produce hormones to signal the stem to reduce elongation. For plants grown in total darkness, the stems continue to grow as if they are still underground to reach the surface before all the nutrition provided by the seed is depleted. Based on our measurements, these processes, however, do not impact the uptake of NaCl or KNO_3_ and hence we found no statistically significant correlation between lighting condition and uptake rate in young tomato seedlings.

### Observing differences in uptake of NaCl in varieties and species of different salt tolerance

Most varieties of *S. lycopresicum* are sensitive to moderate concentrations of NaCl (in the range of hundreds of mM) at various stages of lifecycle, including germination, seedling, and fruit production.^28, 29^ Early phenotyping of behaviors resulting from salt stress, while not able to provide information on fruit yield and adult growth, could be utilized for initial screening of the salt sensitivity of new cultivars. Seedlings are also far more susceptible to high salt levels due to the lack of older tissue, such as leaves, that mature plants use to separate excess salt to protect the rest of the plant.^30, 31^

We investigated the NaCl uptake in both *S. lycopresicum* and a wild species of tomato, *S. pimpinellifolium*, commonly known as the “currant tomato” and known to have a higher salt tolerance compared to the normal tomato.^8^ Six cultivars of *S. lycopresicum* were selected due to their commercial availability: five have had their sensitivities to salt previously established in a study by Gharsallah *et al.* (mildly tolerant: Heinz 1350; sensitive: Marmande, Oxheart, Rio Grande, Saint Peter)^18^; one variety, “Moneymaker”, has been involved in many studies, with conflicting results on the salt sensitivity of the cultivar.^8, 16, 32^ Two varieties of *S. pimpinellifolium* were selected (Golden Currant, Rote Murmel), and although their specific salt tolerances have not been previously studied, *S. pimpinellifolium* is considered to have a higher salt tolerance, with one study suggesting a higher tolerance than the tomato variety “Moneymaker”.^8^

The observed rate of uptake of NaCl (*k*_uptake_) for each variety at the same age (7 days) is shown in **Figure 3A**, against the average dry mass for each variety and their tolerance to NaCl. Although the average dry mass of the two *S. pimpinellifolium* varieties was significantly different than the average dry mass of all the *S. lycopersicum* varieties (**Figure S2A**, one-way ANOVA (*f*(7,16) = 102.7781, *p* < 0.0001)), average dry mass was not shown to be strongly correlated with *k*_uptake_ for the different varieties, with an adjusted r^2^ of only 0.14255 (**Figure S2B**). We did find, however, significant pairwise differences of *k*_uptake_ when comparing different varieties using one-way ANOVA **(Figure 3Bi**, *f*(7,68) = 9.8205, *p* < 0.0001) and a Tukey-Kramer HSD post-hoc test (chosen due to the uneven number of samples of each variety). Both wild tomato varieties showed significant difference in *k*_uptake_ compared to all four of the sensitive varieties of tomato. The sensitive Oxheart variety showed significant difference compared to both the mildly tolerant Heinz 1350 variety and the unknown-tolerance variety Moneymaker. When grouping the different varieties into categories for their sensitivity to NaCl (tolerant, mildly tolerant, sensitive, uncertain), we also found significant difference between the sensitive varieties and the other categories (**Figure 3Bii**, one-way ANOVA (*f*(3,72) = 18.4951, *p* < 0.0001) followed by Tukey-Kramer HSD post-hoc test). No visual changes were observed in the salt-sensitive seedlings over others, but as the amount of salt added was below the concentration usually considered to induce saline stress, this is unsurprising. There were also no visual differences between the different varieties of *S. lycopersicum* before treatment.

**Figure 3.**
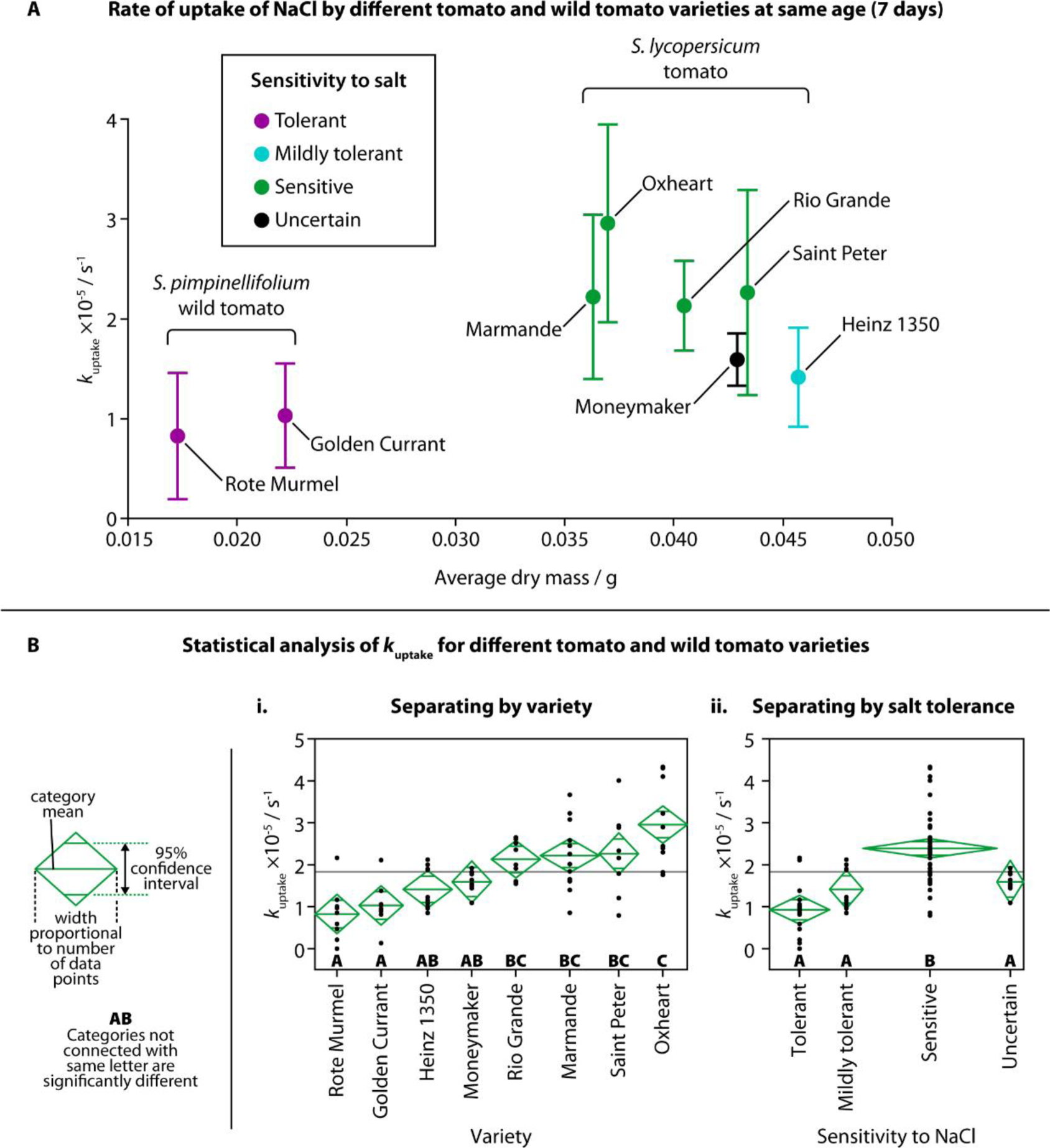
Rate of uptake of NaCl by different tomato and wild tomato varieties. (**A**) *k*_uptake_ for each variety of tomato and wild tomato plotted against the average dry mass for each variety at the same age (7 days). Colors indicate classification of sensitivity to salt (purple = tolerant; cyan = mildly tolerant; green = sensitive; black = uncertain sensitivity). Error bars show one standard deviation (Moneymaker, Saint Peter: n = 8; Golden Currant, Rote Murmel: n = 9; Heinz 1350, Oxheart, Rio Grande: n = 10; Marmande: n = 12). (**B**) Statistical analysis (one-way ANOVA followed by a Tukey-Kramer HSD post-hoc test) showed differences in *k*_uptake_ between some varieties (**i**) and sensitivities to NaCl (**ii**). Confidence diamonds show mean uptake, 95% confidence interval and number of data points, and letters show categories with or without significant difference.

Mechanisms for salt tolerance, Na^+^ uptake and Na^+^ transport are complex, although it is generally agreed that salt-sensitive plants display greater Na^+^ uptake or accumulation.^33^ Transport of Na^+^ from the external root solution to the plant shoot can vary between genotypes.^16^ Na^+^ flow is largely unidirectional, travelling from the roots into the shoot through the relatively fast-moving flow in the xylem. After entering the shoot, Na^+^ is unlikely to return to the roots *via* the phloem, therefore, Na^+^ accumulates elsewhere in the plant, such as leaves.^34^ In some salt resistant species, Na^+^ transport from the roots to the shoot is reduced in the first place – instead, Na^+^ is transported from the basal roots to the distal roots, where Na^+^ is then transported back out into the surrounding solution. Salt sensitive species can also show increased transport of Na^+^ from the roots into the shoot, suggesting higher Na^+^ uptake and upwards transport may be an indicator of salt sensitivity.^20^

We observed higher overall uptake of NaCl in tomato varieties known to be more sensitive to salt, and lower overall uptake by the salt-resistant wild tomato compared to the standard tomato. By showing this relationship, there appears to be a link between NaCl uptake in the seedling stage and the behavior to salt stress in the mature stage. This suggests that TETRIS could prove useful in estimating the salt tolerance of tomato varieties at a very young age, reducing the time taken to test for salt sensitivity by weeks. It is also likely that the wild tomato seedlings exhibited a lower amount of uptake due to their reduced size, as suggested in our earlier experiment comparing different ages of plants; younger plants had both lower fresh mass (Figure S1B) and amount of uptake of NaCl (Figure 2C). The wild tomatoes at an age of 7 days post germination had both a similar fresh mass and amount of uptake of NaCl to Heinz 1350 tomato seedlings at an age of 3 days, as shown in Figure S2C, suggesting both age, mass and genetic makeup play a role in the uptake of salt.

## Conclusions

With this work, we demonstrate our ability to study the effects of physical conditions (*e.g.*, age of plants and light conditions) and biological differences on the uptake rate of ions (NaCl and KNO_3_), using the low-cost soilless electrochemical platform, TETRIS. Our main thesis with the work presented here was whether it is possible to ascertain the salt sensitivity or tolerance of a plant by differences in its uptake of NaCl, in line with physiological traits already studied when looking at salt tolerance.^16^ Not only have we observed significant differences between both species and varieties in uptake, but these differences in rates of uptake of seedlings correspond to the salt sensitivities of the adult plants, suggesting that TETRIS could be used to predict salt resistance. This extrapolation could be useful for screening many hundreds of phenotypes or mutants and picking only the most promising for further experimentation.

The main challenge TETRIS faces is that it currently has only been utilized for young seedlings. Younger plants, however, are more difficult to interface with electrical and electrochemical sensors, due to the fragility of the plant and the weightiness of most sensors, and so TETRIS provides a valuable position for monitoring seedlings. In the majority of experiments, we have grown seedlings on filter paper, an artificial growth environment without sufficient support for older roots. TETRIS consists of low-cost, highly customizable parts made *via* screen-printing and 3D printing and can be easily adapted to different sizes and formats. We have previously shown that TETRIS can be adapted for use in agar, commonly used in plant experiments for *Arabidopsis thaliana*.^23, 35^ An impedance sensor can be embedded into the agar growth medium and monitor as ions are taken up by plants. Theoretically, TETRIS would be compatible with other soilless systems, as long as there is sufficient solution in contact with the electrodes. Soil itself could be more problematic due to the complex and heterogenous geometry, although could be compatible with sufficient coverage of the sensors. The composition of soil includes nutrients and organic matter, considered potential analytes or interferents.^36^ Ion-selective electrodes, such as for K^+^, Ca^2+^ and Na^+^ could be used to decouple the non-specific uptake measured by the impedance sensors in TETRIS and differentiate difference in uptake.^37, 38^

The high customizability and suitability for high-volume manufacturing with our 2D screen-printed sensors allows the use of TETRIS in situations that elude 3D, probe-like electrochemical sensors. Sensors could be embedded into existing soilless growth containers, where the low-profile sensors would not disrupt growth. By applying sensors to the root environment of plants grown in vertical farms, which often use soilless hydroponic systems, TETRIS could be utilized to optimize lighting conditions by measuring differences in nutrient uptake or chemical responses under different light intensities and qualities.^39^ One of the biggest advantages of the screen-printed nature of TETRIS is compatibility with multiplexing, due to the large number of sensors that can be made low-cost (<US $0.10 per sensor). Thousands of samples with differing plant varieties, mutants or growth conditions could be run simultaneously, such as in high-throughput, agar-based optical sensing systems that already exist, where information on the chemical environment would compliment visual differences of the plants.^35^

## Materials and methods

### Impedance sensor fabrication, setup and characterization

Carbon electrodes (Sun Chemical C2130925D1 conductive carbon ink (80 wt%), Gwent Group S60118D3 diluent (20 wt%)) were screen-printed onto polyester transparency sheet (Office Depot). The sensor design consisted of two identical electrodes (30 mm × 6.25 mm) separated by 5 mm. Impedance measurements (amplitude 0.25 V_half-wave_ (RMS), frequency 2 Hz, 0 V d.c.) were performed with a PalmSens 4 potentiostat and MUX8-R2 multiplexers (PalmSens BV, Netherlands). During experiments, the sensor was adhered to the base of a petri dish (55 mm diameter), placed into a measurement chamber consisting of a silicone base and reservoir of water. The sensor was connected to the potentiostat with crocodile clips and a transparent colorless acrylic lid placed over the experimental setup. Paper discs were prepared by printing a hydrophobic wax barrier (HPRT MT800) onto chromatography paper (Whatman, grade 1, 0.18 mm thickness) and heat transferred (Vevor HP230B, 120 °C, 15 minutes) to define a circular area (radius 17.5 mm, area 962 mm^2^). Characterization of NaCl was carried out with 1 ml solution in a paper disc on top of the sensor. The impedance was measured for at least 2 hours and the average of the impedance response recorded.

### Plant growth

The following seeds of *Solanum lycopersicum* (tomato) were obtained: “Cuore Di Bue” (“Oxheart”, Thompson & Morgan UK), “Heinz 1350” (Chiltern Seeds Direct UK, Premiere Seeds Direct UK), “Marmande” (Just Seed UK), “Moneymaker” (Thompson & Morgan UK), “Rio Grande” (Thompson & Morgan UK), “Saint Peter” (“St. Pierre”, Sow Seeds UK). The following seeds of *Solanum pimpinellifolium* (wild tomato) were obtained: “Golden Currant” (Seed-Cooperative UK), “Rote Murmel” (Seed-Cooperative UK). All seeds were stirred in 30% commercial bleach for 15 minutes, rinsed in deionized water and germinated on damp tissue paper for 4 days prior to transfer onto a paper disc. The paper discs with germinated seeds were placed in a propagator box with transparent lid at a relative humidity of around 80%. Ambient light and temperature were used, cycling from around 26°C/22°C, day/night, and 14 hours daylight per day. The paper constantly supplied with deionized water from a reservoir with paper strips.

### Salt uptake experiments

A disc with or without seedlings was removed from the growth chamber after a set number of days (7 days, unless otherwise stated) and placed onto the sensor in the measurement chamber. 100 µL deionized water was added to the paper and, after at least 3 hours, 30 uL salt solution was added to the paper through a port in the lid. To calculate the rate of uptake of the salt, an initial baseline impedance was found by taking the average impedance of the first 2 hours. The time when the impedance was lowest after addition of salt was found and a logarithmic curve of the form log(*Z*) = *t* · *k*_uptake_ + *c* was fitted from this time onwards, where *Z* is impedance, *t* is time, *k*_uptake_ is gradient and the rate of uptake of salt, and *c* is the intercept. Where the curve meets the initial baseline impedance, with a 5% tolerance, the gradient *k*_uptake_ was found for the curve up to that point. Where the line did not reach the initial baseline, a curve was plotted until the end of the experiment time and the gradient *k*_uptake_ was found.

### Drying plants

Plants were removed from the chromatography paper discs and placed into polystyrene weighing boats. They were dried in an oven at 40 °C for at least 16 hours and cooled to room temperature before weighing.

