## Supporting Information for "Plant-on-a-chip: continuous, soilless electrochemical monitoring of salt uptake and tolerance among different genotypes of tomato"

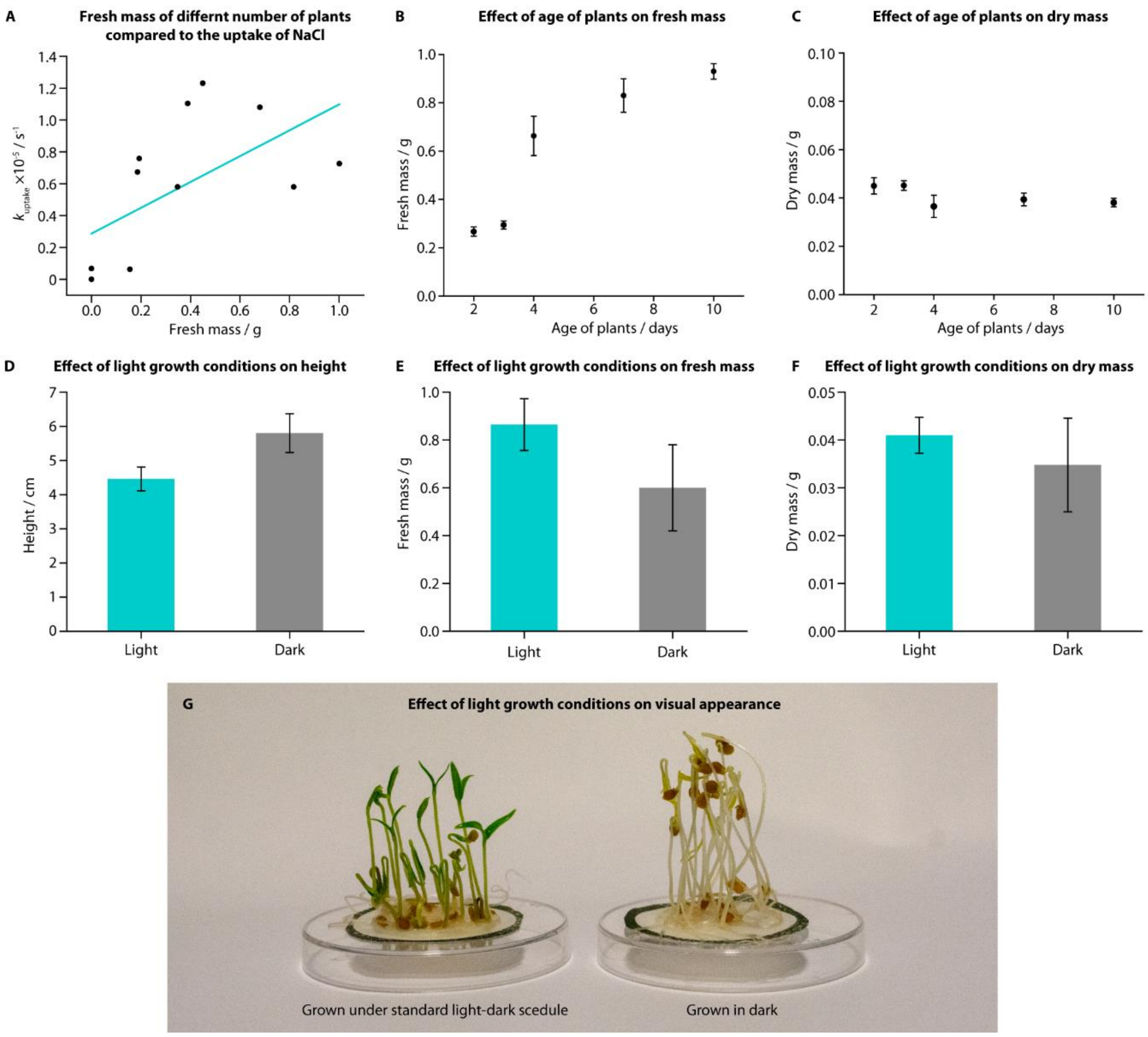


**Figure S1. Uptake of ions, mass and size measured in *S. lycopersicum* "Heinz 1350" under different experimental conditions.** (**A**) $k_{uptake}$ for NaCl (addition of 30 µL, 0.02 M) generally increased with number of tomato seedlings (7 days post germination), which corresponded to the fresh mass of the seedlings after the experiment. (**B**, **C**) Fresh mass of plants generally increased with the age of the plants, but dry mass did not (2 and 3 days: n=6; all others: n = 4). (**D-F**) Although the height of plants grown in the dark was generally greater than plants grown in a standard light-dark schedule, the mass (fresh and dry) of plants grown in the dark was lower. This could be attributed to the health of the plants, which were visibly more yellow and thin (light: n = 5; dark: n = 6). (**G**) Photograph showing the difference between tomato "Heinz 1350" grown under standard light conditions (left) and dark-only light conditions (right) after 7 days growth. Seedlings grown in the dark were noticeably taller and paler.

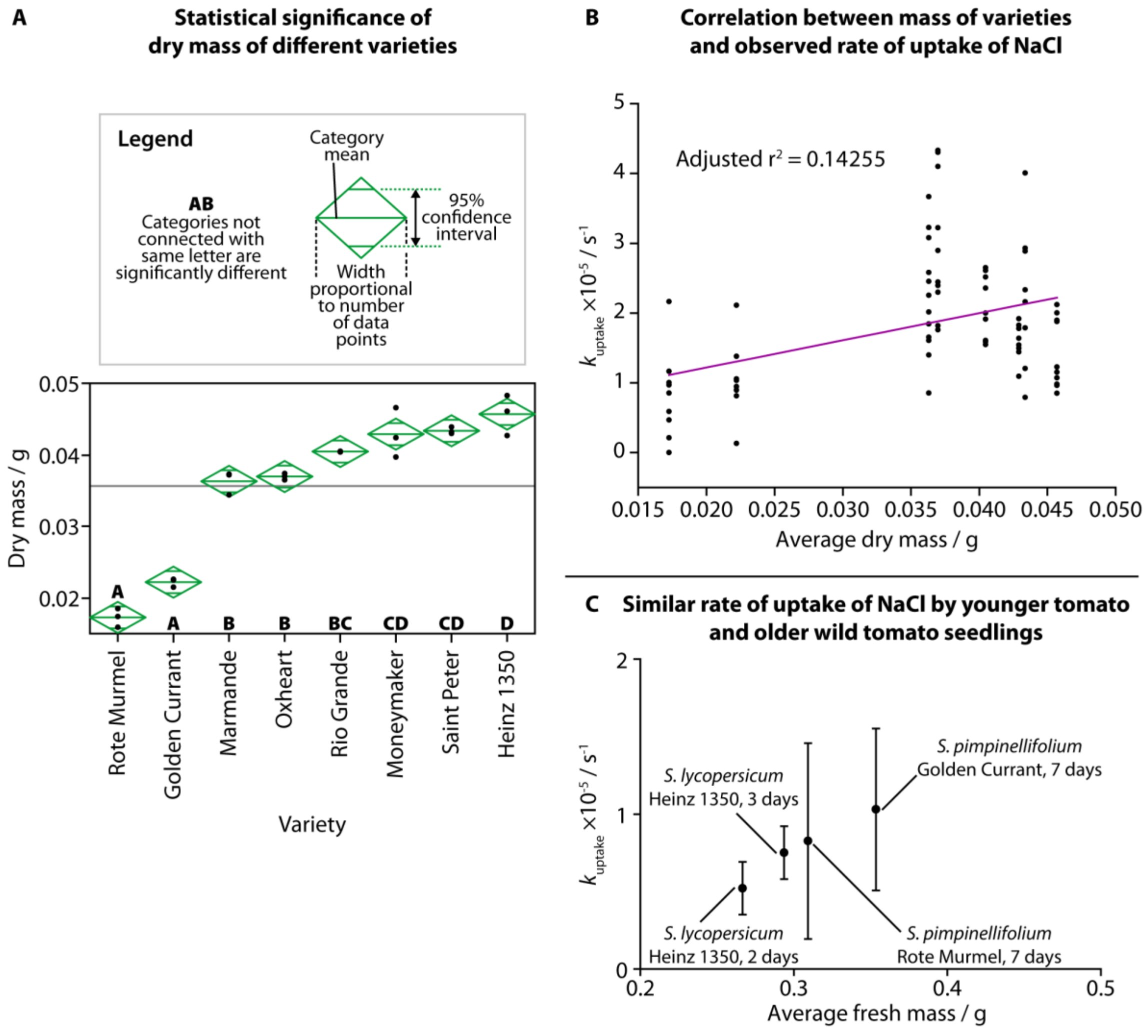


**Figure S2. Mass and uptake of NaCl for different varieties of tomato and wild tomato.** (**A**) Statistical analysis of the dry mass of *S. lycopersicum* and *S. pimpinellifolium* varieties (one-way ANOVA ($f(7,16) = 102.7781$, $p<0.0001$) and Tukey-Kramer HSD post-hoc test). (**B**) Correlation between dry mass and $k_{uptake}$ was weak when comparing all species and varieties from Figure 3A at 7 days old, 20 plants, with a low adjusted $r^2$ of 0.14255. (**C**) Similar rates of uptake of NaCl were observed between younger (2 or 3 days old) tomato plants ("Heinz 1350") and older (7 days old) wild tomato plants, where they also had similar average fresh mass.
